# Gut microbiota and bile acids changes in MASLD mice model with hepatic PLD1 knockout

**DOI:** 10.1101/2025.07.13.663496

**Authors:** Yushang Zhao, Huan Wang, Wanling Lin, Hui Wang, Lin-Lin Cao

**Author notes:** **Correspondence:** Lin-Lin Cao, Department of Clinical Laboratory, Peking University People’s Hospital, #11 Xizhimen South Street, Beijing 100044, China.

## Abstract

Hepatocyte phospholipase D1 (PLD1) knockout alleviated metabolic dysfunction-associated steatotic liver disease (MASLD) in mice, but the underlying mechanism is largely unknown. In this study, high-fat diet is fed to wild type (Con) and hepatocyte PLD1 knockout (Con_KO) mice to establish MASLD model (HFHC and HFHC_KO). Intestinal contents of mice are analyzed via metagenomics and metabolomics, the liver bile acids are assessed by mass spectrometry imaging. At phylum level, Bacillota in the intestines of MASLD model mice are significantly increased and Bacteroidota are significantly decreased. However, after the deletion of hepatocyte PLD1, Pseudomonadota and Candidatus Bathyarchaeota are significantly decreased in the MASLD model mice. Then at species level, compared with Con group, the *Faecalibaculum rodentium* is significantly increased in HFHC group, in which hepatocyte PLD1 knockout causes *Desulfovibrionaceae bacterium* LT0009 and *Lachnospiraceae bacterium* 10-1 to significantly decrease. As for intestinal bile acids, two bile acids (Hyodeoxycholic acid and Glycolithocholic acid) are found to be different between the HFHC_KO group and the HFHC group. Association analysis shows the *Faecalibaculum* co-occurs with DCA, βMCA, ΩMCA and αMCA, while probiotic *Bacteroides uniformis* is significantly correlated with UDCA, 12-KetoLCA, 7-KetoLCA. Finally, mass spectrometry imaging reveals that TCA and TDCA in liver are significantly decreased after hepatocyte PLD1 knockout. These findings demonstrate that hepatocyte PLD1 knockout alters gut microbiota and bile acids profiles, suggesting PLD1 deficiency may modulate MASLD progression by changing intestinal microbiota-bile acid homeostasis.

**Highlights:** Here, we show that hepatocyte PLD1 knockout alters gut microbiota and bile acid profiles in metabolic fatty liver disease mouse by high-fat diet.

1. Wild type (Con) and hepatocyte PLD1 knockout (Con_KO) mice were used to establish HFHC and HFHC_KO models, respectively.
2. Intestinal contents were collected for metagenomic and metabolomics analysis, and liver tissues were taken for mass spectrometry imaging to investigate gut microbiota-bile acid relationships.
3. In HFHC_KO mice, *Desulfovibrionaceae bacterium* LT0009 and *Lachnospiraceae bacterium* 10-1 were significantly reduced, accompanied by altered HDCA and GLCA.
4. Association analysis revealed *Faecalibaculum* co-occurred with DCA, βMCA, ΩMCA, and αMCA, while *Bacteroides uniformis* was significantly associated with UDCA, 12-KetoLCA, and 7-KetoLCA.
5. Mass spectrometry imaging showed hepatocyte PLD1 knockout significantly decreased liver TCA and TDCA, suggesting PLD1 deficiency may modulates MASLD progression via microbiota-bile acid homeostasis.

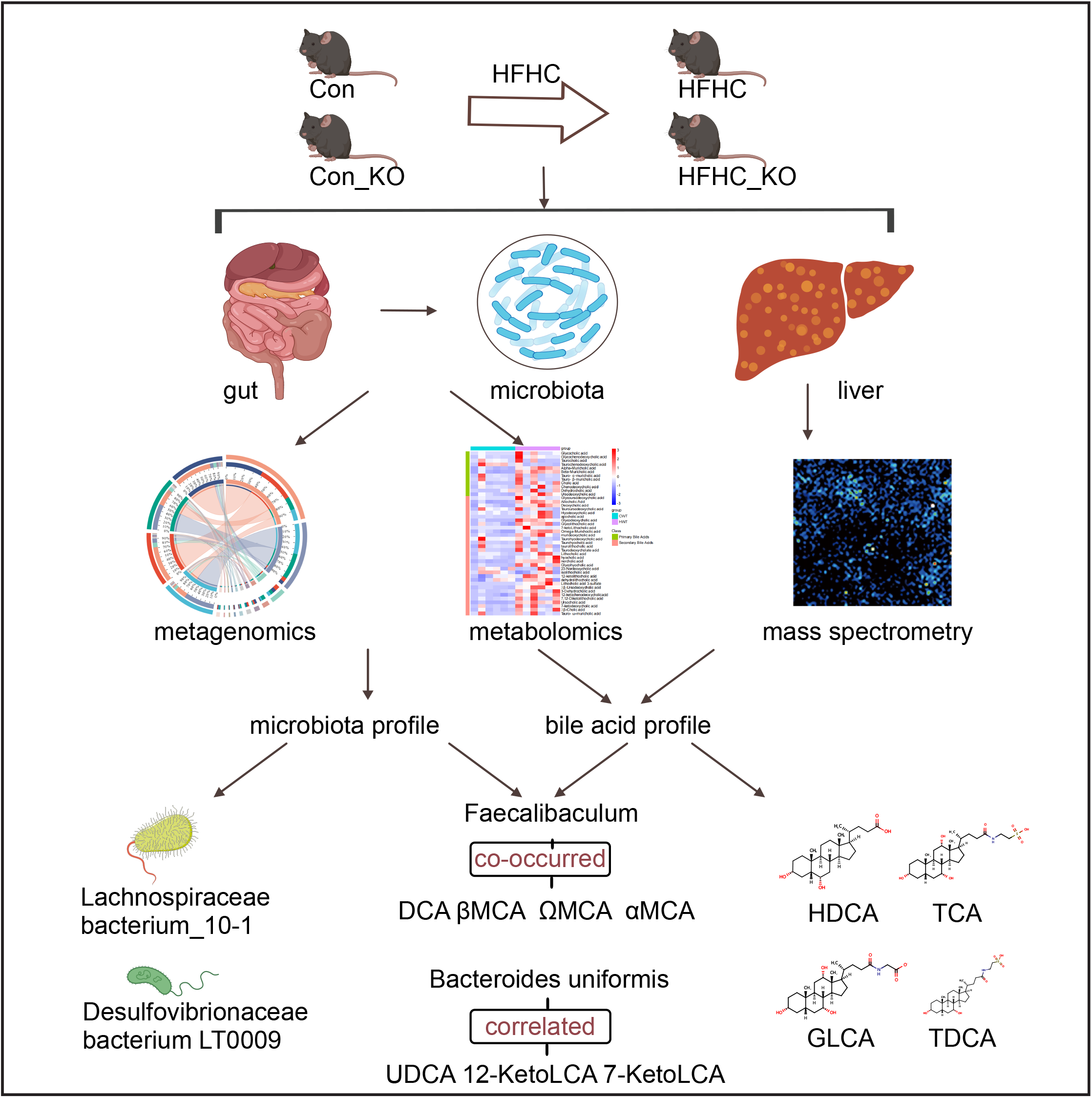

## Introduction

MASLD has emerged as the most prevalent chronic liver disease, with the transformation of modern lifestyle and the increasing prevalence of metabolic syndrome such as obesity, the prevalence of MASLD is likely to continue rising. MASLD is associated with an increased risk of cardiovascular events, extrahepatic malignancy, liver failure, and hepatocellular carcinoma[1-3]. Nevertheless, to date, there remains a lack of clear and reliable therapeutic approaches for MASLD in clinical practice.

An increasing number of researchers are now exploring the interaction between the gut-liver axis and MASLD, using gut microbiota as a key entry point to search for interventions for liver lesions in MASLD. Previous studies have demonstrated that the structure of gut microbiota in MASLD patients often undergoes significant alterations, and the most typical feature is the imbalance in the ratio of Bacillota to Bacteroidetes, which is closely associated to the development of hepatic steatosis and obesity, this strongly indicates the potential role of dysbiosis of the gut microbiota in the pathogenesis of MASLD[4]. In patients with metabolic dysfunction-associated steatohepatitis (MASH), the abundance of *Bacteroides* and *Ruminococcus* was significantly increased and *Prevotella* was decreased[5]. In addition, animal studies have provided compelling evidence for the potential role of gut microbiota in MASLD[6]. Studies have found that the ingestion of intestinal microbiota from MASLD mice by normal mice can cause the transmission of MASLD-related disease symptoms[7].

Phospholipase D1 (PLD1), an isoenzyme of phospholipase D (PLD), has a unique biological function, which can decompose phosphatidylcholine into phosphatidic acid. As an important lipid second messenger, phosphatidic acid is widely involved in the regulation of intracellular signaling pathways[8]. PLD1 is ubiquitously expressed on the surface of mammalian cells[8], and participates in lipid metabolism by regulating the accumulation of lipid droplets[9, 10]. In our prior studies, it was found that PLD1 was significantly up-regulated in the liver of MASLD mice, and the disease were markedly alleviated when hepatic PLD1 was absent[11]. Although previous studies have highlighted the close relationship between gut microbiota and MASLD, there is currently no evidence regarding the impact of hepatic PLD1 on gut microbiota.

In this study, we investigated the effects of hepatic PLD1 on gut microbiota and bile acid metabolism in a high-fat diet induced MASLD mouse, and discussed whether this could explain the remission of MASH disease caused by hepatic PLD1 knockout. We conducted a series of rigorous experiments. Specifically, intestinal contents of mice were subjected to metagenomic and untargeted metabolomics sequencing, while liver tissue was subjected to bile acid metabolomic mass spectrometry imaging analysis. The results showed that in MASLD mice, Bacillota was significantly increased and Bacteroidetes was significantly decreased. However, when hepatic PLD1 was absent, Pseudomonadota and Candidatus Bathyarchaeota was significantly decreased. Moreover, corresponding changes in bile acids were observed, and a certain correlation was found between the microbiota and bile acids.

## Materials and methods

### 1. Animal model

Hepatocyte PLD1 knockout mice were generated at the Shanghai Model Organisms Center, Inc (Shanghai, China). All animals were housed in a pathogen-free, temperature-regulated facility at Beijing Friendship Hospital, which followed a 12-hour light/dark cycle. All animal procedures were approved by the Institutional Animal Care and Ethics Committee. For the experiment, 8–9-week-old hepatocyte PLD1 knockout mice and wild-type mice were provided with either a normal control diet or a high-fat diet rich in fat and cholesterol (western diet composition: 50% sugar, 20% protein, 21% fat, 1% cholesterol; sourced from Research Diets, New Brunswick, New Jersey, USA). After 16 weeks of feeding, mouse feces were collected in a sterile environment, and then these mice were sacrificed to obtain livers.

### 2. Metagenomic analysis

0.5g of stool was used to extract total genomic DNA with the FastPure Stool DNA Isolation Kit(Magnetic bead) (MJYH, shanghai, China) according to manufacturer’s instructions. The concentration and purity of the extracted DNA were assessed using SynergyHTX and NanoDrop2000, respectively. DNA quality was verified via 1% agarose gel electrophoresis. For paired-end library construction, the DNA extract was fragmented to an average size of approximately 400 bp using a Covaris M220 (Gene Company Limited, China). The paired-end library was generated with the NEXTFLEX Rapid DNA-Seq kit (Bioo Scientific, Austin, TX, USA). Sequencing was conducted on an Illumina NovaSeq™ X Plus (Illumina Inc., San Diego, CA, USA) at Majorbio Bio-Pharm Technology Co., Ltd. (Shanghai, China), employing the NovaSeq X Series 25B Reagent Kit following the manufacturer’s protocols (www.illumina.com). Metagenomic sequencing data were analyzed on the Majorbio Cloud Platform (www.majorbio.com).

### 3. Bile acid metabolomic analysis

The 47 bile acids standards were weighed accurately and prepared as single standard master mixes in methanol. The working standard solution was prepared by measuring an appropriate amount of each master batch and diluting it with 50% acetonitrile to a suitable concentration. The isotope standards (CDCA-D4, CA-D4) were weighed accurately and prepared as single master batch with methanol. The working standard solution was added to the internal standard solution at a volume ratio of 1:1 and mixed well to obtain differnt concentration solutions.

Add 50 μL of internal standard working solution (200 ng/mL) and 350 μL of extraction solution (methanol), vortex and mix for 30 s, sonicate at low temperature for 30 min (5°C, 40 KHz), than store at -20°C for 30 min, centrifuge at 4°C for 15 min at 13000 rcf, remove the supernatant and blow dry with nitrogen, add 100 μL of 50% acetonitrile, vortex for 30 s, sonicate at 5L (40 KHz) for 10 min, then centrifuge at 4°C for 15 min at 13000 rcf, collect the supernatant for LC-MS/MS. The supernatant was purged with nitrogen, added 100 μL of 50% acetonitrile, vortexed for 30 s, sonicated at 5 □ (40 KHz) for 10 min, centrifuged at 4 □ for 15 min at 13000 rcf, and the supernatant was used for LC-MS/MS analysis.

The LC-MS/MS analysis was performed on an ExionLC AD system coupled with a QTRAP® 6500+ mass spectrometer (Sciex, Boston, USA) at Majorbio Bio-Pharm Technology (Shanghai, China). Samples were separated using a Waters BEH C18 column (150×2.1 mm, 1.7 μm) maintained at 50 □. The mobile phases consisted of solvent A (water with 0.025% formic acid and 10 mM ammonium acetate) and solvent B (acetonitrile:methanol, 9:1 v/v), delivered at 0.4 mL/min. The gradient program was: 0–0.5 min, 5%–10% B; 0.5–1.0 min, 10%–20% B; 1.0–2.0 min, 20%–28% B; 2.0–8.0 min, isocratic at 28% B; 8.0–11.0 min, 28%–33% B; 11.0–20.0 min, 33%–45% B; 20.0–23.0 min, 45%–50% B; 23.0–29.0 min, 50%–80% B; 29.0–29.01 min, 80%– 100% B; 29.01–31.0 min, isocratic at 100% B; 31.0–31.01 min, 100%–5% B; 31.01–33.0 min, isocratic at 5% B for system equilibration. Samples were stored at 4 □ during analysis. Mass spectrometric data were collected using an ESI source in negative ion mode on a SCIEX QTRAP 6500+ with optimized parameters: CUR 35, CAD Medium, IS-4500 V, TEM 550 □, GS1 50, GS2 50. Raw data were imported into Sciex OS software, and metabolite concentrations were calculated via linear regression standard curves.

### 4. mass spectrometry imaging

The mass spectrometry imaging was carried out by Shimadzu. The chemicals involved were from Sigma-Aldrich(St.Louis, MO, USA): α-cyano-4-hydroxycinnamic acid (CHCA, 97%), 1,5-diaminonaphthalene (DAN, 97%), Girard’s reagent T (GIT, 98%). The preparation of the liver section was performed according to a normal procedure. Livers were sectioned at -20 ° C and mounted on ITO slides (1200nm conductive layer thickness, South China Technology Co., LTD.). Finally, the sections were subjected to laser irradiation.

For the analysis of cinnamaldehyde and methoxy-cinnamaldehyde with air pressure MALDI mass spectrometry imaging (AP-MALDI MSI), they were firstly derivatized with GIT and then applied with CHCA as matrix. GIT and CHCA were prepared at 10.0 mg/ml in 20% acetic acid and 50% acetonitrile aqueous solution (0.1% TFA), respectively. Leave GIT derivatization for 80min at room temperature and then spray CHCA onto sample surface. Both GIT and CHCA were sprayed with an iMlayer AERO automatic sprayer (Shimadzu, Kyoto, Japan). The common spraying parameters were as follows: layers, 20; pumping pressure: 0.1MPa; spraying pressure: 0.2MPa; drying time, 1s; washing frequency, 1 layer/wash; scan pitch, 1mm. The flow rate, nozzle distance, and stage speed were different: 0.015 and 0.06 ml/min, 9 and 5 cm, 120 and 70 mm/s for GIT and CHCA, respectively. For the rest compounds, they were detected in negative ion mode with DAN as matrix. DAN was vapor-deposited onto sample surface with an iMLayer™ matrix vapor deposition system (Shimadzu, Kyoto, Japan). Deposition parameters were as follows: temperature, 180 □; vacuum, 2×10-2Pa; time, 5min.

Mass spectrometry imaging was performed on iMScope QT (Shimadzu, Kyoto, Japan) equipped with an optical microscope, an AP-MALDI source, and a quadrupole time-of-flight mass spectrometer (LCMS-9030). Statistical analysis and image reconstruction were carried out with IMAGEREVEAL™ MS (Shimadzu, Kyoto, Japan). Normalization was performed for each sample with TIC as denominator. Minimum threshold value was set as 0% to avoid missing any possible active ingredients. The average value of weighted mass peak area of each region of interest (ROI, generally the whole sample) was used as indicator of relative content for each active ingredient.

### 5. Statistical analysis

Statistical analysis was performed using GraphPad Prism software (version 10.1.2, San Diego, California, USA), and values were expressed as mean ± SEM. Differences between two groups were compared using Student’s t test for normal distribution and the Mann–Whitney test for abnormal distribution. Statistical significance was set at p <0.05.

## Results

### 1. High-fat diet significantly altered the diversity of intestinal microbiota in mice

In our previous study, it was found that hepatocyte PLD1 knockout could mitigate high-fat diet induced MASLD in mice. In order to investigate the impact of hepatocyte PLD1 knockout on intestinal microbiota and bile acid metabolism, hepatocyte PLD1 knockout mice and normal control mice were randomly assigned to two groups and fed either a high-fat diet or a normal control diet. The mice were divided into four groups: Con (normal control diet), HFHC (high-fat diet), Con_KO (hepatocyte PLD1 knockout with normal control diet), and HFHC_KO (hepatocyte PLD1 knockout with high-fat diet). Metagenomic sequencing of the intestinal microbiota was performed after we collected intestinal contents from mice. The results demonstrated that the differences across groups were generally greater than the differences within groups (Figure 1A). Mice fed a high-fat diet exhibited significantly different intestinal microbiota distribution characteristics compared to those fed a normal diet, while minimal differences were observed between the Con group and Con_KO group, as well as between the HFHC group and HFHC_KO group (Figure 1B, C). In terms of microbial diversity, the Shannon index and Chao index of intestinal microbiota in HFHC group and HFHC_KO group were significantly lower than those in the other two groups, respectively, while there was no significant difference between HFHC group and HFHC_KO group (Figure 1D). Further analysis of the gut microbial distribution at phyla level showed that Bacillota and Bacteroidota were predominant in all groups (Figure 1F). Therefore, High-fat diet had a more pronounced effect on the diversity of gut microbiota than did hepatic PLD1 in mice.

**Figure 1.**
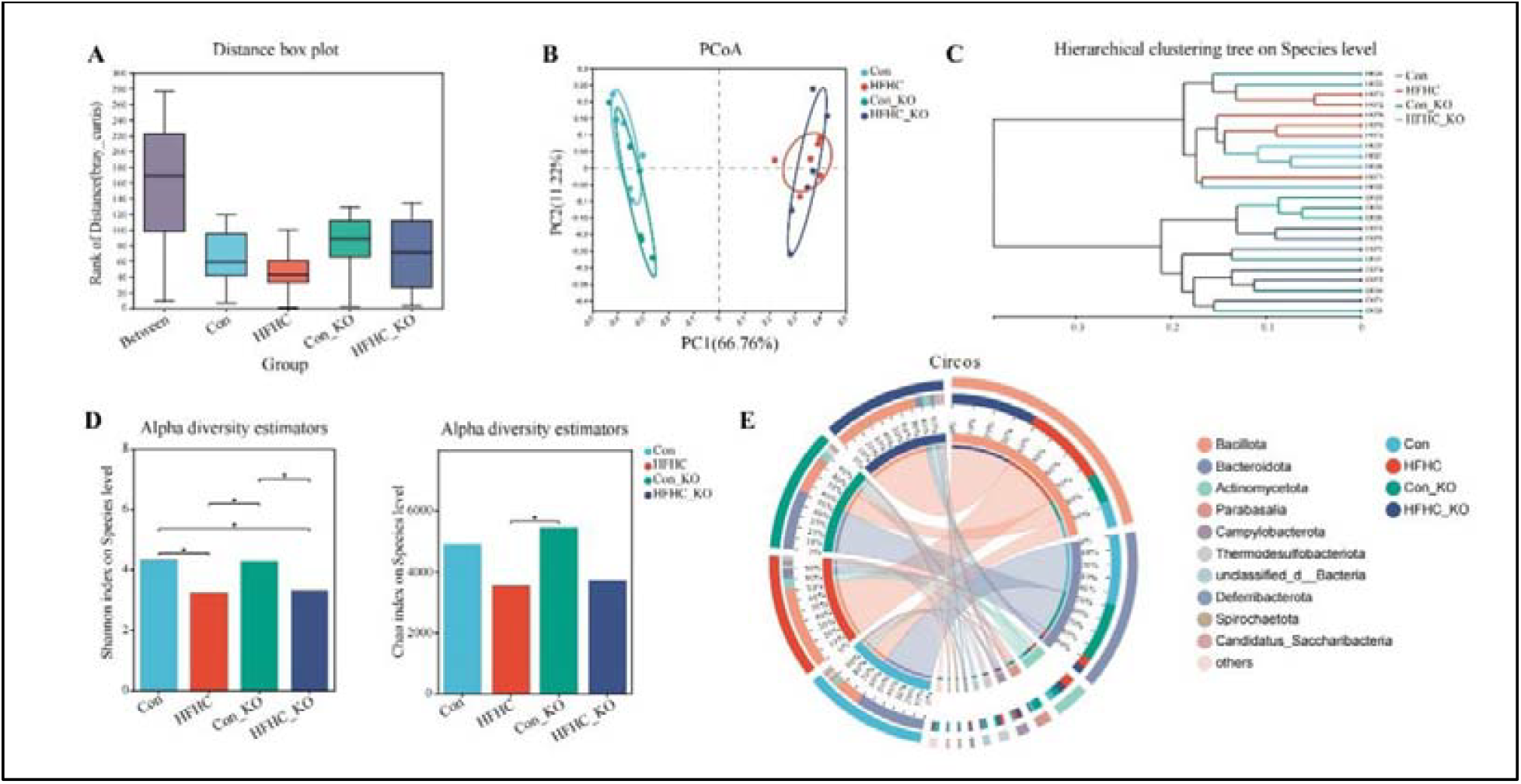
High-fat diet changed the diversity of intestinal flora in mice. Overall status of intestinal microbiota diversity in each group of mice. (A) The box plot was used to show the differences between groups and between groups, The ordinate represents the distance value. The boxes “Between” refer to differences between groups, while the others represent differences within each group. (B) The similarity and difference of community between groups were demonstrated by using two-dimensional visual scatter plots, and the degree of agglomeration and dispersion of sample was reflected by the distance between samples. (C) The sample-level clustering tree was used to show the similarities and differences of species composition among samples. (D) Difference of Chao index and Shannon index among different groups. (D) Circos sample-species diagrams was used to show the distribution of species present in different group. On one side of the graph are the groups, and on the other side are the dominant species. Data are presented as mean ± SEM, n = 6 mice per group. ^*^p <0.05.

### 2. Hepatocyte PLD1 knockout led to significant alterations in the abundance of certain bacteria species

At the phylum level, compared to the mice on a normal diet, those fed a high-fat diet exhibited a notable increase in the abundance of Bacillota and a significant decrease in Bacteroidota. However, when comparing the HFHC group with the HFHC_KO group, no significant differences were observed in these two dominant phyla above (Figure 2A, C). Additionally, the levels of Pseudomonadota and Candidatus Bathyarchaeota were significantly lower in the HFHC_KO group than in the HFHC group (Figure 2B). Further analysis at the species level revealed that HFHC mice exhibited a significant increase in *Faecalibaculum rodentium* compared to Con mice, along with notable changes in other low-abundance species. Meanwhile, compared to the HFHC group, HFHC_KO mice showed significant decreases in *Desulfovibrionaceae bacterium* LT0009 and *Lachnospiraceae bacterium* 10-1 (Figure 2D, E). By intersecting the differentially changed species between the Con group and the HFHC group, as well as between the HFHC group and the HFHC_KO group, a total of 18 differentially expressed species were identified. Among these 18 species, *Desulfovibrionaceae bacterium* LT0009 and *Lachnospiraceae bacterium* 10-1 had the highest abundance (Figure 2F, Table 1). Previous studies have indicated that *Lachnospiraceae* can produce long chain fatty acids (elaidate) by decomposing high-fat foods, and elaidate can damage the intestinal barrier and cause metabolic disorders such as obesity. Although it has little impact on diversity, hepatic PLD1 knockout has indeed led to significant changes in the abundance of *Desulfovibrionaceae bacterium* LT0009 and *Lachnospiraceae bacterium* 10-1.

**Figure 2.**
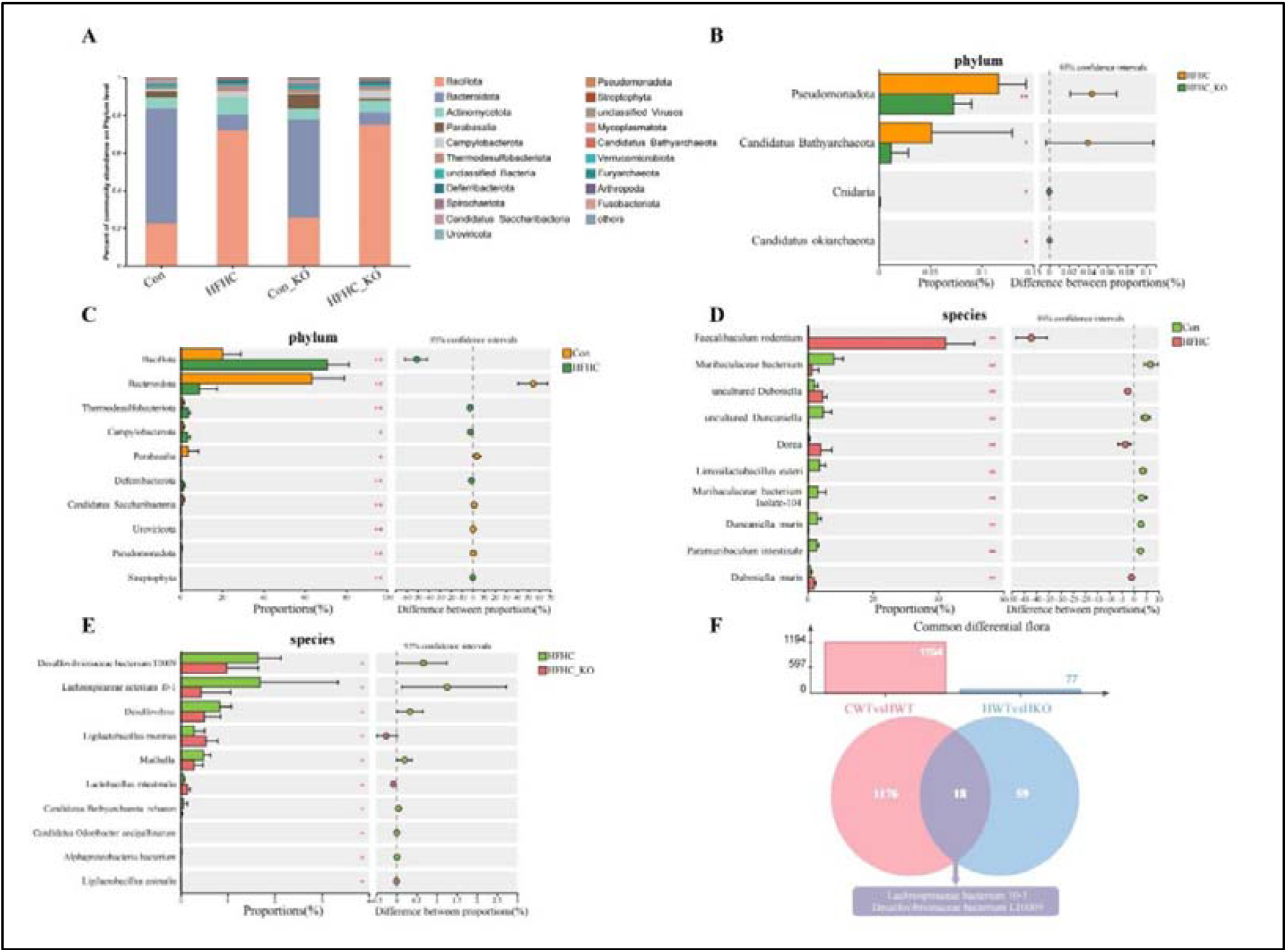
Liver PLD1 knockout resulted in a significant change in the abundance of a subset of bacteria. (A) The community bar chart showed the species composition of the top 20 abundant in all samples and the proportion of different species, and other low-abundance species are classified into others. (B) The bar chart showed the difference in mean relative abundance of the different phylum between HFHC group and HFHC_KO group. (C) The bar chart showed the difference in mean relative abundance of the different phylum between Con group and HFHC group. (D) The bar chart showed the difference in mean relative abundance of the different species between Con group and HFHC group. (E) The bar chart showed the difference in mean relative abundance of the different species between HFHC group and HFHC_KO group. (F) 18 species were obtained by intersection of Con vs HFHC mice and HFHC vs HFHC_KO mice. Data are presented as mean ± SEM, n = 6 mice per group. ^*^p <0.05, ^**^p <0.01.

**Table 1.**
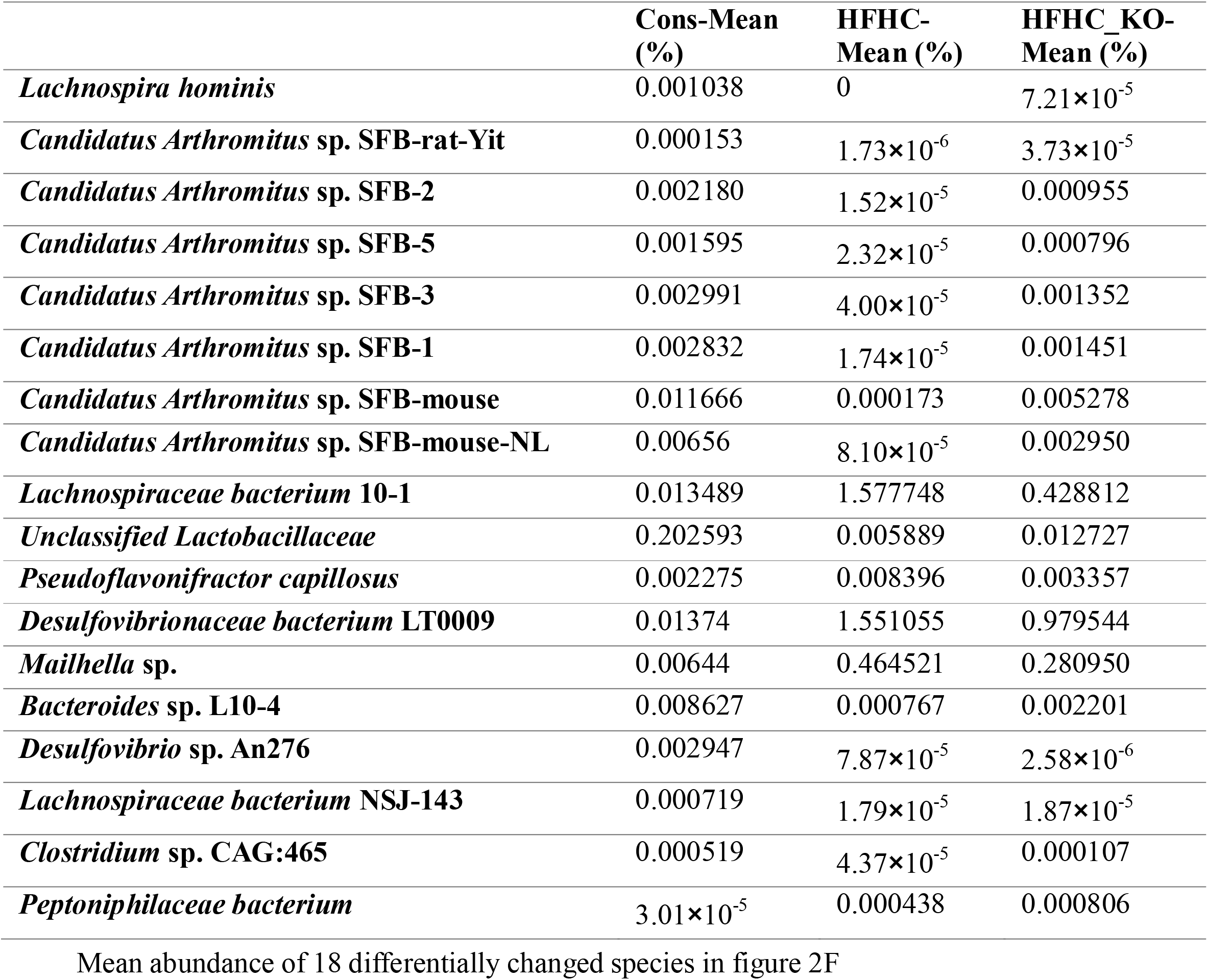
Mean Abundance.

|  | **Cons-Mean (%)** | **HFHC-Mean (%)** | **HFHC_KO-Mean (%)** |
| --- | --- | --- | --- |
| *Lachnospira hominis* | 0.001038 | 0 | $7.21 \times 10^{-5}$ |
| *Candidatus Arthromitus* sp. SFB-rat-Yit | 0.000153 | $1.73 \times 10^{-6}$ | $3.73 \times 10^{-5}$ |
| *Candidatus Arthromitus* sp. SFB-2 | 0.002180 | $1.52 \times 10^{-5}$ | 0.000955 |
| *Candidatus Arthromitus* sp. SFB-5 | 0.001595 | $2.32 \times 10^{-5}$ | 0.000796 |
| *Candidatus Arthromitus* sp. SFB-3 | 0.002991 | $4.00 \times 10^{-5}$ | 0.001352 |
| *Candidatus Arthromitus* sp. SFB-1 | 0.002832 | $1.74 \times 10^{-5}$ | 0.001451 |
| *Candidatus Arthromitus* sp. SFB-mouse | 0.011666 | 0.000173 | 0.005278 |
| *Candidatus Arthromitus* sp. SFB-mouse-NL | 0.00656 | $8.10 \times 10^{-5}$ | 0.002950 |
| *Lachnospiraceae bacterium* 10-1 | 0.013489 | 1.577748 | 0.428812 |
| *Unclassified Lactobacillaceae* | 0.202593 | 0.005889 | 0.012727 |
| *Pseudoflavonifractor capillosus* | 0.002275 | 0.008396 | 0.003357 |
| *Desulfovibrionaceae bacterium* LT0009 | 0.01374 | 1.551055 | 0.979544 |
| *Mailhella* sp. | 0.00644 | 0.464521 | 0.280950 |
| *Bacteroides* sp. L10-4 | 0.008627 | 0.000767 | 0.002201 |
| *Desulfovibrio* sp. An276 | 0.002947 | $7.87 \times 10^{-5}$ | $2.58 \times 10^{-6}$ |
| *Lachnospiraceae bacterium* NSJ-143 | 0.000719 | $1.79 \times 10^{-5}$ | $1.87 \times 10^{-5}$ |
| *Clostridium* sp. CAG:465 | 0.000519 | $4.37 \times 10^{-5}$ | 0.000107 |
| *Peptoniphilaceae bacterium* | $3.01 \times 10^{-5}$ | 0.000438 | 0.000806 |

Mean abundance of 18 differentially changed species in figure 2F

### 3. Hepatocyte PLD1 knockout changed the gut microbiome metabolism of mice

Initially, our findings revealed that the intestinal microorganisms across all groups of mice were predominantly enriched within metabolic pathways (Figure 3A, B). Cluster analysis of the enriched metabolic pathways revealed that the metabolic environment in the gut of mice on a high-fat diet was markedly different from that of mice on a normal control diet, while the effect of gene knockout was relatively subtle (Figure 3C). Further investigation through bile acids metabolomics sequencing analysis of intestinal contents demonstrated that when comparing the HFHC group with the Con group, a total of 23 bile acids exhibited significant changes. In contrast, only two bile acids, namely hyodeoxycholic acid (HDCA) and glycinocholic acid (GLCA), underwent significant alterations when the HFHC_KO group was compared with the HFHC group. (Figure 3D,E). Intriguingly, these two bile acids did not coincide with the previously mentioned 23 bile acids. Remarkably, we observed a high degree of consistency between the metagenomic data and the bile acid metabolome data (Figure 3F). Collectively, at the metabolic level, the hepatic knockout of PLD1 significantly modified the composition of bile acids in the gut, with HDCA and GLCA being prime examples of such alterations.

**Figure 3.**
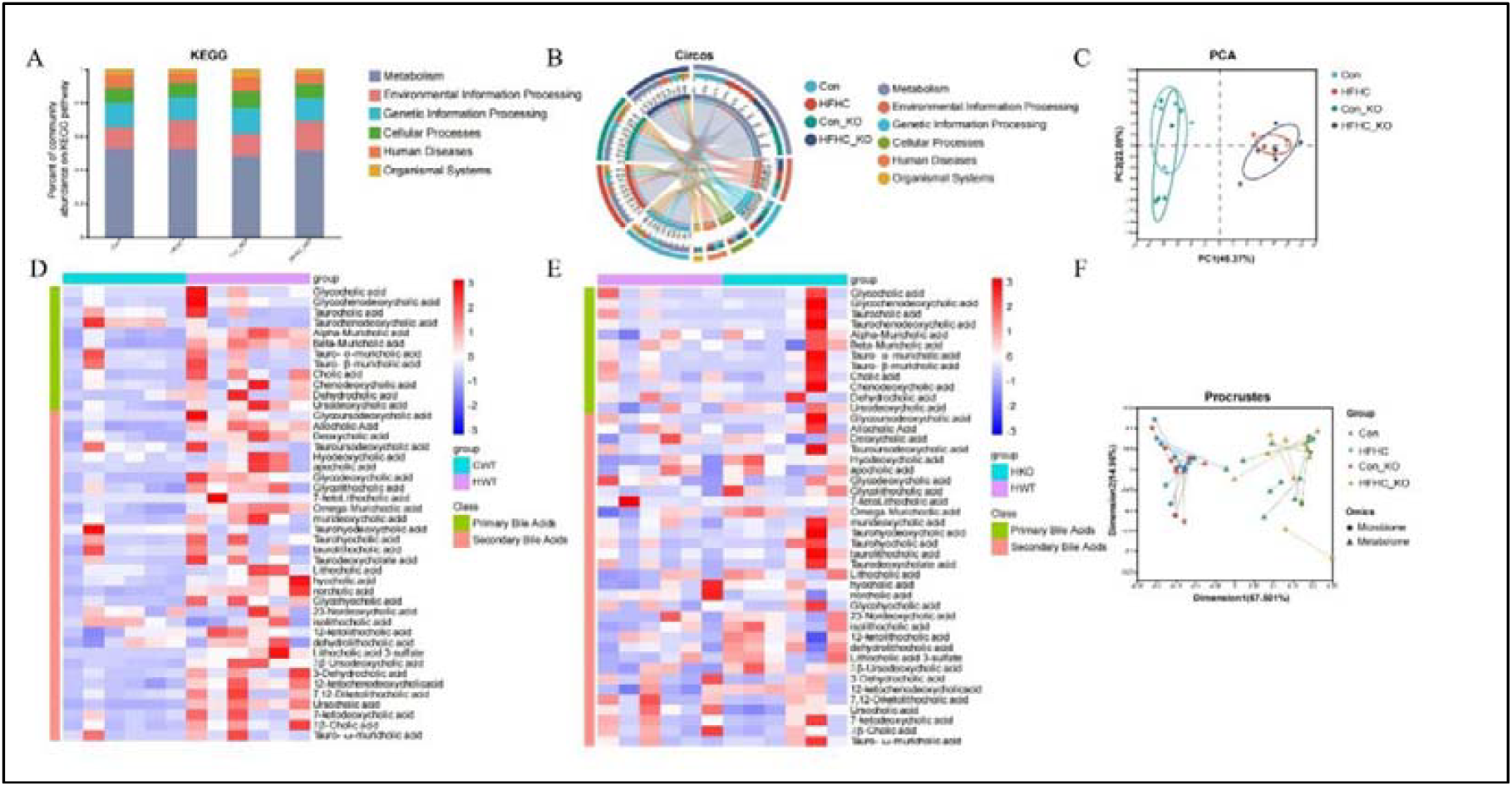
The western diet significantly changed the gut microbiome metabolism of mice. (A)The KEGG Bar chart showed the composition of KEGG functions and the proportion of different functions in each group. (B) Circos sample and function diagram was used to show the distribution of KEGG function present in different group. On one side of the graph are the samples and their groups, and on the other side are the KEGG function items. (C) Scatter plot showed the similarity and difference of the functions of different samples, and reflects the degree of convergence and dispersion of the functions of samples by the distance between samples (Euclidian distance algorithm). (D, E) Heat map showed the distribution of bile acids in different groups of samples. (F) Procrustes analysis plot shows the consistency of samples across different data sets. Each line segment represents a sample, and the lines represent the Procrustes residuals of the two ordering configurations. The shorter the lines, the higher the consistency between the two data sets. n = 6 mice per group.

### 4. Bile acids in gut and the liver were correlated with bacterial abundance in the gut

To further explore these relationships, we performed co-occurrence probability analysis with the top 50 bacteria genera and all 46 measured bile acids, and the findings revealed that DCA, βMCA, ΩMCA, and αMCA exhibited a particularly high tendency to co-occurred with all bacteria genera. Among the bacteria, *Faecalibaculum* showed the maximum co-occurrence probability with all four bile acids (Figure 4A). Previous studies have confirmed that *Bacteroides uniformis* exerts a beneficial effect on liver metabolic diseases. In the present study, *B. uniformis* is significantly correlated with UDCA, 12-KetoLCA and 7-KetoLCA bile acids, However, it is likely that other bacteria species may also be involved in the metabolism of these bile acids (Figure 4BCDE). In addition, we also employed mass spectrometry imaging to analyze liver tissues from four groups of mice. The results indicated that the contents of TCA and TDCA in the liver of the HFHC group were significantly increased compared with the Con group, while the contents of TCA and TDCA in the HFHC_KO group were significantly decreased compared with the HFHC group (Figure 4FGH). This discrepancy from the changes observed in intestinal contents may be attributed to the enterohepatic circulation and the distinct microenvironmental characteristics of the liver and intestine. In conclusion, our findings demonstrate a correlation between bile acids and the abundance of microbiota, although the causal relationship remains to be elucidated, these results suggest a potential mechanism by which phospholipase D1 (PLD1) may influence bile acid metabolism through its effects on the gut microbiota.

**Figure 4.**
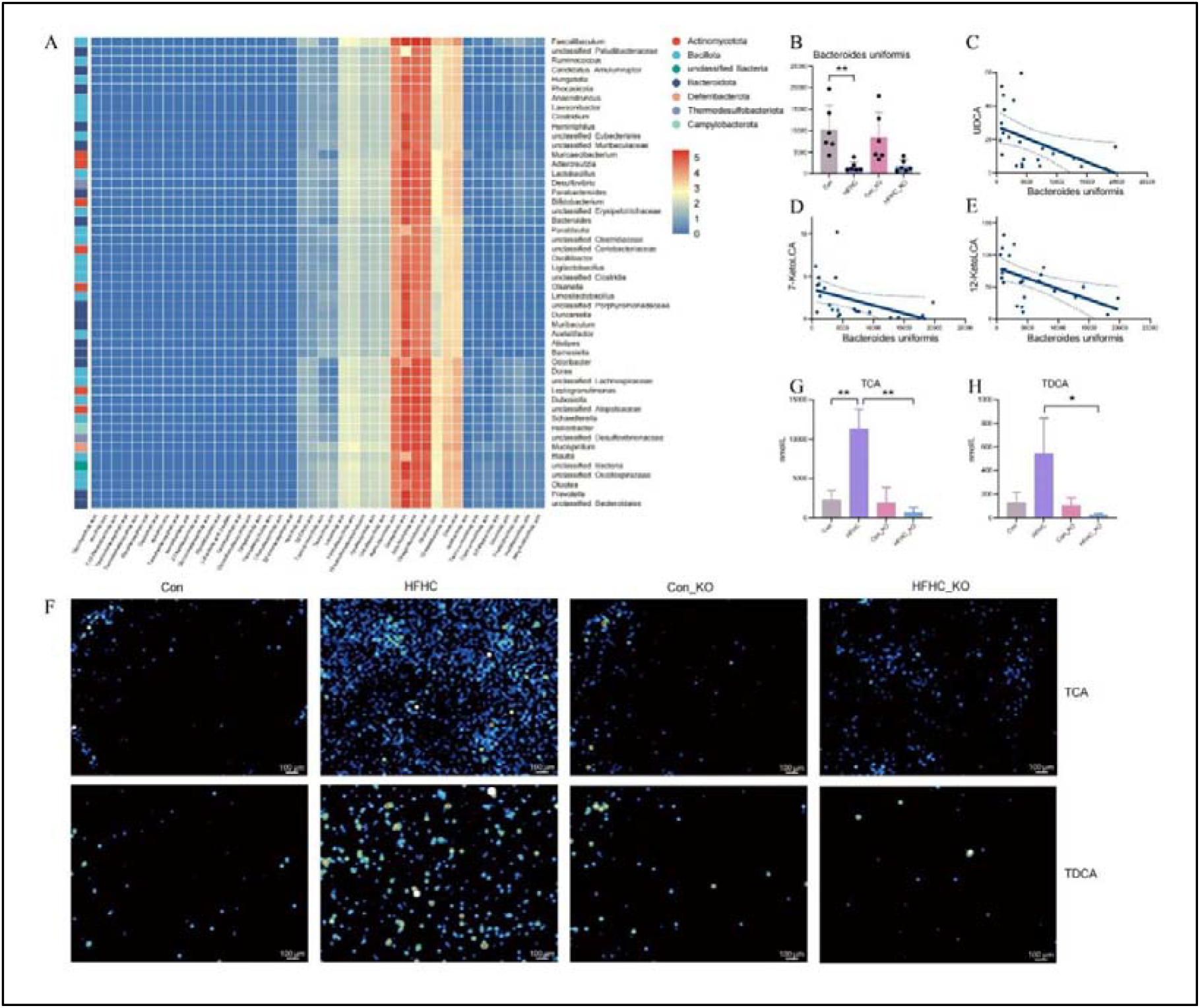
Bile acids in intestinal contents and the liver were correlated with bacterial abundance in the gut. (A) Mmvec analysis diagram showed co-occurrence probabilities between microorganisms and bile acids. Microorganisms with top50 co-occurrence probability were selected for plotting. The color bar shows the logarithmic conditional probability between microorganisms and bile acids, the redder the color, the greater the probability of their co-occurrence. (B) The histogram shows the abundances of Bacteroides uniformis in different groups. (C, D, E) Correlation plot showed the correlation between bile acids and Bacteroides uniformis. (F) The mass spectrogram showed the contents of TCA and TDCA in different groups. (G, H) The histogram showed the contents of TCA and TDCA in different groups according the F diagram. Data are presented as mean ± SEM, n = 6 mice per group. ^*^p <0.05, ^**^p <0.01.

## Discussion

Upon exposure to a high-fat diet, the gut microbiota composition and bile acid profiles undergo profound transformations. In previous investigations, remarkable shifts in intestinal microbe were consistently observed in MASLD disease. Animal studies have provided compelling evidence: inoculating normal mice with the gut microbiota from MASLD mice can trigger the manifestation of MASLD, strongly suggesting a potential causal link between the gut microbiota and the pathogenesis of MASLD[12]. Additionally, several research endeavors have unearthed a positive correlation between *Bacteroides* or *Ruminococcus* and the severity of MASLD in human patients[5]. While species-specific disparities between mouse and human gut microbiota pose challenges for translational research, fecal microbiota transplantation (FMT) has emerged as a valuable tool. FMT from patients with MASH into germ-free mice has successfully recapitulated key features of MASH, including hepatic steatosis and inflammation. These pathological changes are further exacerbated by high-fat diet consumption[13].

In the present study, metagenomic analysis revealed that high-fat diet intervention elicited profound changes in the gut microbiota composition of mice, such as a decrease in Bacteroidota and an increase in Bacillota, in which *Faecalibaculum rodentium* increased. Intriguingly, liver-specific knockout of PLD1 induced less dramatic changes to the gut microbiota compared to high-fat diet exposure, particularly in terms of microbiota diversity. This phenomenon may be attributed to the fact that the genetic alterations only affect hepatic cells, exerting limited effect on the intestinal environment.

Notably, we found two species which changed significantly: *Lachnospiraceae bacterium* 10-1 and *Desulfovibrionaceae bacterium* LT0009. However, due to the restriction of bacterial isolation method, their characteristics and effects were not further studied. It has been reported that *Lachnospiraceae* can act as a probiotic to protect the intestinal barrier through its metabolite butyrate, thereby improving the body and liver metabolism[14-16]. However, other researchers have found that the commensal species *Fusimonas intestini* can damage the intestinal barrier by decomposing high-fat foods to produce long-chain fatty acids—Elaidate, causing metabolic disorders such as obesity [17]. As for *Desulfovibrio*, the *Desulfovibrio* genus has been shown to be dominate when a high-fat diet impairs gut microbiota composition and is associated with cancer progression[18].

Given that PLD1 gene editing occurs exclusively in the liver and does not directly affect the gut microbiota, the question arises: what mediates the connection between the two? We suspect that the most likely mediator is bile acids, since the enterohepatic circulation of bile acids is the most direct way for the gut to interact with the liver. Indeed, bile acids play a crucial role in the enterohepatic circulation in MASLD. Bile acids generally exhibit a certain toxic effect on bacteria, yet under specific circumstances, they can promote microbial diversity. Emerging evidence from multiple studies indicates that the microbiota diversity in patients with cholestasis is reduced[19], and in mice, bile duct ligation leads to a decrease in microbial β-diversity[20]. Although the underlying mechanism has not been fully elucidated, it is hypothesized that bile acids secretion may serve as an energy substrate to support the survival and proliferation of multiple microorganisms, thereby contributing to the maintenance of microbial diversity.[21]. TUDCA mitigate disease progression in MASLD mice by reducing intestinal inflammation, restoring intestinal barrier integrity, reducing intestinal fat transport and regulating intestinal microbiota composition[22]. As a newly discovered probiotic, *B. uniformis* reduces MASH by producing a modified bile acid 3-sucCA, which promotes the growth of *Akkermansia muciniphila*[23].

Therefore, we performed bile acid metabolomics analysis of intestinal contents. We found that dietary conditions had a strong effect on bile acids, but we did not identified any bile acids that could directly explain the phenomenon of “ PLD1 knockout alleviated fatty liver induced by high-fat diet”. We found that *Faecalibaculum* was significantly co-dominant with bile acids including DCA, αMCA, βMCA and ΩMCA, However, the causal interplay among them remains undefined. Therefore, our subsequent research will employ strategies such as microbiota transplantation to systematically investigate the bidirectional relationship between bile acid profiles and the gut microbial community. This approach aims to investigate whether *Faecalibaculum* modulates bile acid metabolism or if these bile acids shape *Faecalibaculum* abundance, shedding light on the mechanistic linkages underlying their co-dominance. Besides, no significant correlations were found between bile acids and the two differentially abundant bacteria, *Desulfovibrionaceae bacterium* LT0009 and *Lachnospiraceae bacterium* 10-1. In addition, previous studies have demonstrated that *B. uniformis* exert beneficial effects on liver metabolic diseases such as MASLD through *A. muciniphila* [23], and we found that it was significantly increased in our MASLD mouse model, and it was significantly correlated with three beneficial bile acids, UDCA, 12-KetoLCA, and 7-KetoLCA. However, these correlations were not positive. We suggested that other bacteria may also contributes to bile acid metabolism. And the increase of *B. uniformis* may be an adaptive response of the intestinal ecosystem in order to counteract the pathological damage.

Notably, our findings reveal a discrepancy in bile acid profiles between the intestine and liver, which is likely attributed to the different metabolic processes and regulatory mechanisms of bile acids in the intestines and the liver. In the liver, cholesterol is converted into primary bile acids through a series of enzymatic reactions initiated by the rate-limiting enzyme CYP7A1[24]. Conversely, In the intestine, the presence of gut microbiota drives the transformation of primary bile acids into secondary bile acids via bacterial bile salt hydrolase (BSH) and 7α-hydroxylase activities. FXR is highly expressed in the gastrointestinal tract, serves as a critical bile acid sensor. It mediates feedback inhibition of hepatic bile acid synthesis, thereby maintaining exceedingly low bile acid concentrations in hepatocytes to prevent liver injury and cholestasis. Notably, Different bile acids have significantly different efficacy in activating FXR, further influencing metabolic outcomes [24]. Collectively, these integrated mechanisms—including organ-specific metabolic transformations and FXR-mediated regulatory loops—give rise to the marked divergence in bile acid compositions between the liver and intestine.

If the intestinal microbiota alterations induced by PLD1 knockout are attributed to changes in bile acids, how exactly do these bile acid modifications occur? We postulate two potential mechanisms. Firstly, PLD1 in the liver may regulate the expression of bile acid synthesis enzymes, such as CYP7A1 and CYP8B1, which are crucial for the biosynthesis of primary bile acids. Secondly, the knockout of PLD1 may alter the activity of microbial enzymes involved in bile acid metabolism, especially the 7α-hydroxylase, which is necessary for the formation of secondary bile acids. To clarify which mechanism is operative, we plan to employ PCR analyses to simultaneously quantify the expression of key bile acid metabolism enzymes in liver tissues and the intestinal microbiota after liver PLD1 knockout. This method will help elucidate whether PLD1 directly regulates bile acid metabolism through the liver enzymatic pathway or indirectly regulates bile acid metabolism through microbial metabolic activities.

Additionally, as this study is fundamentally a correlational one, it has certain limitations in exploring functions and mechanisms. In the subsequent research, the phenomenon can be further confirmed by overexpressing PLD1. Besides, in vitro studies using organoids can also be carried out to delve into the underlying mechanisms. Whereas, it should be noted that the animal model employed in this study was mice, whose microbiota composition exhibits substantial differences from that of humans. For further research on human MASLD, two potential approaches could be adopted: transplanting human intestinal contents into mice or conducting direct investigations in human patients.

In conclusion, the gut microbiota and bile acid showed significant changes after high-fat diet feeding. Hepatic PLD1 knockout may alter the composition of bile acid by disturbing intestinal microbiota homeostasis, and then influence the disease progression of MASLD.

## Acknowledgement

We thank Zhenhe Chen and Xiaojuan Lei in Shimadzu for mass spectrometry imaging.

